# Reconstitution of glycan-driven MHC I recycling reveals calreticulin as mediator between TAPBPR and tapasin

**DOI:** 10.1101/2025.11.25.690393

**Authors:** Tim J. Heinke, Amin Fahim, Niko Popovic, Tobias Rath, Nina Morgner, Simon Trowitzsch, Robert Tampé

## Abstract

Protein folding in the endoplasmic reticulum (ER) is essential for about one-third of the mammalian proteome. N-linked glycosylation and subsequent glycan remodeling barcodes glycoproteins during their maturation in the ER. Major histocompatibility complex class I (MHC I) molecules, key for adaptive immunity, rely on a dedicated quality control cycle that involves specialized chaperones and glycan-modifying enzymes for their maturation and loading of immunogenic peptides. However, the functional interplay of the MHC I editors tapasin as part of the peptide-loading complex (PLC), TAP-binding protein-related (TAPBPR), the UDP-glucose:glycoprotein glucosyltransferase 1 (UGGT1), and calreticulin in glycan-dependent transfer of MHC I clients has not been determined in a reconstituted system. With isolated components, we show that transfer of peptide-receptive MHC I from the downstream quality control factor TAPBPR back to tapasin depends on the recognition of the monoglucosylated glycan of MHC I by calreticulin. While calreticulin’s C-terminal acidic helix is dispensable for disengaging reglucosylated MHC I from TAPBPR, it is essential for docking MHC I onto tapasin. Our data provide a mechanistic basis for glycan-surveillance by calreticulin necessary for retrograde trafficking of misfolded or suboptimally loaded MHC I that escaped the first quality control at the PLC and were trapped by TAPBPR. Such finetuned dynamic network of glycan-dependent and MHC I-specific chaperones guarantees maturation of MHC I molecules and highlight the fundamental processes driving ER protein quality control.

**Teaser:** Our study dissects the mechanistic network of dedicated chaperones and glycan modifiers that quality-control MHC I.

**Significance Statement:** The immune system relies on major histocompatibility complex (MHC) class I molecules to present protein fragments from within cells, enabling detection of infection or disease. This study uncovers how a dynamic, glycan-dependent chaperone network in the endoplasmic reticulum (ER) orchestrates the recycling of misfolded or suboptimally loaded MHC I. Using isolated components, the work shows how monoglucosylation of MHC I glycans by UDP-glucose:glycoprotein glucosyltransferase 1 (UGGT1) allows calreticulin to mediate transfer from the post-ER chaperone TAPBPR back to the ER-resident chaperone tapasin as part of the peptide-loading complex. These findings illuminate the coordinated action of TAPBPR, UGGT1, calreticulin, and tapasin-ERp57 in MHC I quality control, offering new insights in immune surveillance and how its disruption may contribute to disease.

## Introduction

Protein folding in the endoplasmic reticulum (ER) is crucial for 30–40% of the mammalian proteome comprising membrane-associated, secreted, and organelle-targeted proteins (1, 2). A cleavable N-terminal ER-targeting signal peptide emerging from nascent chains at the ribosome allows proteins to be translocated across the ER membrane and to enter the secretory pathway (3, 4). Most translocated proteins are N-linked glycosylated by the oligosaccharyltransferase (OST), which transfers a Glc_3_Man_9_GlcNAc_2_ appendage from a dolichol phosphate precursor that resides in the ER membrane (5). OST scans the emerging polypeptide for glycosylation sequons Asn-X-Ser/Thr and adds the preassembled oligosaccharide to the asparagine side chain (4, 6).

Transferred N-linked glycans are rapidly restructured by ER-resident glycosidases and transferases whereby the composition of the glycan acts as a dynamic barcode of the maturation status of the protein it is attached to (7, 8). Deglucosylation occurs in a controlled sequential manner and is initiated by α-glucosidase I that removes the outermost α-1,2-linked glucose (5, 9). The remaining α-1,3-linked glucose moieties of the Glc_2_Man_9_GlcNAc_2_ glycan are stepwise removed by α-glucosidase II, licensing client proteins to leave the ER and to enter the secretory pathway (5, 10). The specialized carbohydrate-binding chaperones calreticulin (Crt) and calnexin specifically bind monoglucosylated Glc_1_Man_9_GlcNAc_2_ glycans on client proteins and thereby control trafficking of proteins in the secretory pathway (11).

As another crucial determinant in ER protein quality control, UDP-glucose:glycoprotein glucosyltransferase 1 (UGGT1) regulates glycoprotein folding and degradation (12) by adding back a terminal glucose to Man_9_GlcNAc_2_ glycans of misfolded or misassembled glycoproteins, allowing an additional round of chaperone-mediated folding in the calnexin/calreticulin cycle (11). UGGT1 recognizes non-native or near-native glycoproteins with exposed hydrophobic regions (13–15) using a central, hydrophobic cavity in its protein-sensing domain (16–18). As type I membrane glycoproteins, MHC I heavy chains (hc) in complex with β_2_-microglobulin (β_2_m) are also subject to quality control by the calnexin/calreticulin cycle (19–21). Furthermore, UGGT1-mediated quality control is crucial for maturation of MHC I molecules (22, 23) and is promoted by the MHC I-specific chaperone TAPBPR (24, 25).

MHC I molecules present peptide ligands, derived from proteasomal degradation of proteins in the cytosol, on the cell surface of all nucleated cells and are key for the adaptive immunity to detect and eliminate infected and diseased cells (26–28). The ATP-binding cassette (ABC) transporter associated with antigen processing (TAP1/2) shuttles these peptide ligands into the ER lumen (20, 29), where they undergo further trimming by aminopeptidases ERAP1 and ERAP2 generating peptides of optimal length for fitting in the MHC I binding groove (30–33). Initial loading and selection of high-affinity peptides is coordinated by the peptide-loading complex (PLC), a highly dynamic supramolecular machinery in the ER, consisting of TAP1/2, the MHC I-dedicated chaperone tapasin, the protein disulfide isomerase ERp57, as well as Crt and MHC I molecules (19, 34). Tapasin and TAPBPR stabilize intrinsically unstable peptide-deficient MHC I, and accelerate peptide exchange and selection of high-affinity peptide ligands (19–21, 35–45). Although both MHC I-dedicated chaperones share 20% sequence identity, have similar interfaces to MHC I, and fulfill similar functions (46–49), tapasin primarily acts in the peptide-rich environment of the ER as a key constituent of the PLC, whereas TAPBPR does not associate with the PLC and instead acts in the peptide-depleted ER/Golgi compartment (24, 40, 42, 50).

Crt is a conserved Ca^2+^-binding chaperone that is predominantly localized in the ER lumen (51–53). Crt possesses an N-terminal globular lectin-like domain responsible for glycan recognition, a central proline-rich (P) domain, and a highly acidic C-terminal region involved in Ca^2+^-buffering, followed by an ER-retention motif (Lys-Asp-Glu-Leu, KDEL) (51). Various Crt mutants that impede its chaperone or Ca^2+^-buffer capacities have been implicated in malignant transformation, tumor progression, and response to cancer therapy (54, 55). Such Crt mutants often show changes in their C-terminal domain bearing a novel, positively charged amino acid sequence and lacking the KDEL motif and thus enabling mutant Crt to escape the ER. Whereas Crt uses its lectin domain to recruit MHC I via the monoglucosylated glycan at Asn86 of their heavy chain (hc) (56), it uses the tip of its P domain to bind to residues in the b’ domain of ERp57 (57–59) and its C-terminal acidic helix to latch onto the C-terminal ER-lumenal domain of tapasin (34, 60). The coordinated interplay between Crt binding sites on MHC I molecules, tapasin, and ERp57 is crucial for MHC I assembly, maturation in the PLC, and antigen processing (53, 61). Whether a glycan-independent binding component is involved in Crt-mediated recruitment of MHC I to tapasin-ERp57 within the PLC remains unclear to date.

Although quality control of MHC I molecules is primarily mediated via the PLC involving tapasin and Crt and subsequently by UGGT1 and TAPBPR (24, 25), it is not known how MHC I molecules that did not pass the second quality control are shuttled back to the PLC. Furthermore, the mechanisms that are executed by Crt for quality control of peptide-MHC I complexes in the secretory pathway are currently not fully understood (62). The sequence and intermediates governing MHC I transitions between TAPBPR, UGGT1, Crt, and tapasin-ERp57 remain incompletely resolved. Existing cellular studies lack the temporal and biochemical control necessary to delineate these transient and sequential processes, motivating us to develop a fully reconstituted and compositionally defined in-vitro system. Here, we set out to characterize teamwork between the two MHC I-dedicated chaperone systems to systematically investigate MHC I-chaperone exchange reactions under defined conditions. Our detailed mechanistic study reveals the crucial role of monoglucosylation of MHC I N-glycans and the specific contribution of Crt and its acidic C-terminal helix in the sequential transfer of peptide-receptive MHC I molecules from TAPBPR to tapasin-ERp57. We also show that MHC I retained and reglucosylated by UGGT1/TAPBPR is shuttled back to tapasin-ERp57 in a Crt-dependent fashion. We demonstrate that reglucosylation by UGGT1 is essential for Crt to capture MHC I from a TAPBPR-chaperoned state. Additionally, we show that the C-terminal acidic helix of Crt is dispensable to detach peptide-receptive MHC I from TAPBPR but is essential for recruiting peptide-receptive MHC I to tapasin-ERp57. Our findings reveal the mechanistic basis of a fine-tuned MHC I chaperone handover and network of facilitators which is crucial for an effective immune response.

## Results

### Engineering glycan-driven MHC I chaperone handover in vitro

The ER folding environment is highly complex: Client proteins encounter multiple chaperones and glycan-modifying enzymes in dynamic equilibria that pose a considerable challenge for mechanistic studies *in cellulo*. To overcome this, we aimed to design a fully reconstituted in-vitro system using purified components to precisely control the glycosylation state of MHC I molecules and to sequentially monitor their interactions with TAPBPR, Crt, and the tapasin-ERp57 complex. We selected the TAPBPR-dependent, yet tapasin-independent, MHC I allomorph HLA-A*68:02 (63, 64) to enable robust detection of chaperone exchange events. To analyze UGGT1-dependent transfer of MHC I from the secondary chaperone TAPBPR to the primary MHC I chaperone tapasin in vitro (Fig. 1 *A*), we first co-expressed and secreted a fusion construct comprising the ectodomain of HLA-A*68:02 hc and β_2_m (termed MHC I hereafter), equipped with a cleavable C-terminal Fos leucine zipper motif. Additionally, we included the ER-lumenal domain of TAPBPR with a cleavable C-terminal Jun leucine zipper motif and a Twin-Strep-tag (25, 65). To establish soluble TAPBPR/MHC I heterodimers with a defined mannose-only Man_9_GlcNAc_2_ glycan attached to Asn86 of the MHC I hc, we took advantage of the α-mannosidase I inhibitor kifunensine (66, 67) during expression and secretion of the zippered constructs from HEK293 cells (25). We isolated the secreted zippered MHC I-TAPBPR complex by affinity chromatography, removed the Jun-Fos zipper motifs by Tobacco Etch Virus (TEV) protease treatment, and purified the complex by size exclusion chromatography (SEC) (*SI Appendix*, Fig. S1 *A*). We next produced recombinant tapasin-ERp57 heterodimer, comprising the ER-lumenal domain of tapasin and a stabilizing Cys36Ala mutant of full-length ERp57 using insect cell expression (48) (*SI Appendix*, Fig. S1 *B*). To separate TAPBPR-containing from tapasin-containing complexes by SEC, we increased the hydrodynamic radius of the tapasin-ERp57 heterodimer by attaching a Fab fragment of the well-described PaSta1 antibody (68) to tapasin (*SI Appendix*, Fig. S1 *B*). The PaSta1 Fab binds on top of the N-IgV domain of tapasin without affecting its interactions with ERp57 or MHC I (49). Lastly, we isolated recombinant His-tagged Crt or Crt lacking parts of its C-terminal acidic helix (termed Crt^Δacidic^ hereafter) from *E. coli* extracts by immobilized-metal affinity chromatography (IMAC) followed by SEC (*SI Appendix*, Fig. S1 *C*). We next assayed the chaperone handover of MHC I from TAPBPR to tapasin by assessing complex formation under competitive conditions using SEC. The glycan status of MHC I was determined by liquid chromatography coupled mass spectrometry (LC-MS) (*SI Appendix*, Table S1).

**Fig. 1.**
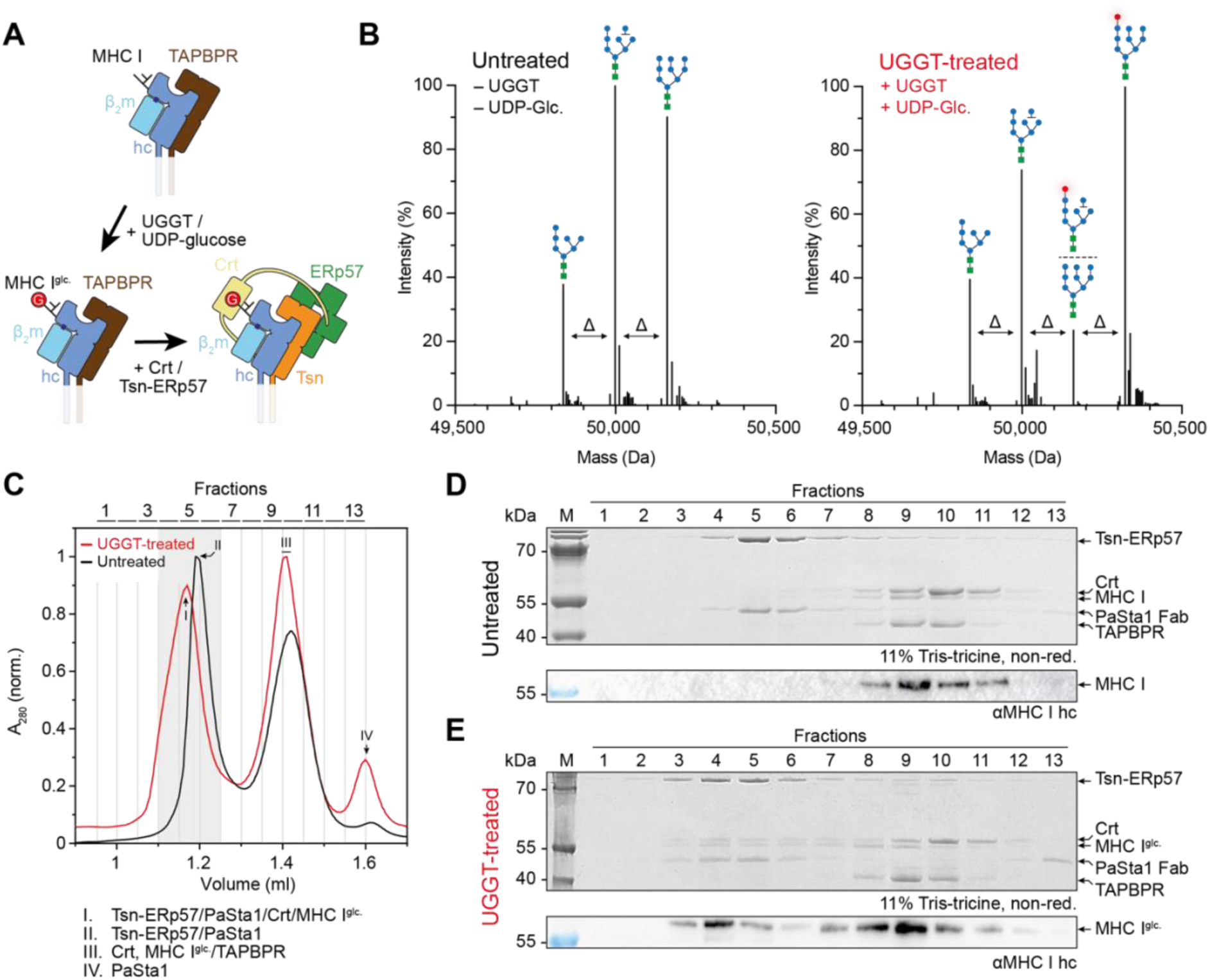
Transfer of MHC I from TAPBPR to tapasin dependent on monoglucosylated glycan. (*A*) Schematic representation of the handover assay, highlighting the interaction between MHC I heavy chain (hc, blue), β_2_-microglobulin (β_2_m, cyan), TAPBPR (brown), calreticulin (Crt, yellow), ERp57 (green), and tapasin (Tsn). The glucose moiety of the monoglucosylated glycan (Glc_1_Man_9_GlcNAc_2_) of the MHC I heavy chain (hc) is marked by a red-circled G. (*B*) Deconvoluted mass spectra of single-chain β_2_m-HLA-A*68:02 displaying the native glycan pattern (untreated) and UGGT/UDP-glucose-treated single-chain β_2_m-HLA-A*68:02 (UGGT-treated). A mass shift of 162 Da (Δ), corresponding to one hexose moiety, is indicated (Δ). Peak labels show the structure of the N-glycan attached to the β_2_m-HLA-A*68:02 single chain construct (green dot – GlcNAc, blue dot – Man, red dot – Glc). (*C*) Size exclusion chromatography (SEC) analyses of β_2_m-HLA-A*68:02 containing either an unmodified Man_9_GlcNAc_2_ glycan (untreated, black) or a monoglucosylated Glc_1_Man_9_GlcNAc_2_ glycan (UGGT-treated, red) in the presence of tapasin-ERp57-PaSta1, TAPBPR, and Crt (3 µM each). Grey region indicates the differences in migration range of tapasin-ERp57-PaSta1 and MHC I-Crt-tapasin-ERp57-PaSta1 complexes. Roman numbers above each peak relate to proteins or protein complexes defined below. (*D,E*) Non-reducing SDS-PAGE and immunoblotting (using the HC10 antibody against MHC I heavy chain) of SEC fractions for untreated (*D*) or UGGT-treated (*E*) single-chain β_2_m-HLA-A*68:02. M – marker, I – input, kDa – kilodalton, A_280_ – absorption at 280 nm, αMHC I – anti-MHC I heavy chain. Data in panels *C-E* are representative of three independent experiments (n = 3).

### A monoglucosylated glycan of MHC I is essential for chaperone transfer

We first studied glycan dependency of the MHC I transfer from TAPBPR to tapasin by our SEC-based approach (Fig. 1 *A*). To this end, we incubated TAPBPR-chaperoned peptide-receptive MHC I with *Chaetomium thermophilum* (*Ct*) UGGT in the presence of UDP-glucose and Ca^2+^ ions (25). We used *Ct*UGGT throughout the study (termed UGGT hereafter) as it generates the same glycan pattern as human UGGT1 but shows better stability and faster reaction kinetics. Purified MHC I-TAPBPR heterodimers were specifically monoglucosylated by UGGT, producing an MHC I species predominantly carrying the Glu_1_Man_9_GlcNAc_2_ monoglucosylated N-glycan (termed MHC I^glc.^ hereafter) as confirmed by LC-MS (Fig. 1 *B* and *SI Appendix*, Table S1). UGGT also monoglucosylated glycans missing one mannose at either the B or C branch, resulting in glycan species of the Glu_1_Man_8_GlcNAc_2_ type, although with lower efficiency (Fig. 1 *B* and *SI Appendix*, Table S1). The lower reglucosylation efficiency of Man_8_GlcNAc_2_ compared to Man_9_GlcNAc_2_ glycan is consistent with the previously described activity of human UGGT1 (69). UGGT preferentially monoglucosylates Man_9_ over Man_8_ or Man_7_ glycoforms, recognizing the innermost GlcNAc residue of the Man_9_GlcNAc_2_ glycan (70, 71). When we compared the handover of TAPBPR-chaperoned monoglucosylated and mannose-only MHC I to tapasin in the presence of Crt using SEC, we observed the formation of a kinetically stable complex between MHC I, Crt, tapasin-ERp57, and PaSta1 only when the MHC I hc carried the monoglucosylated Glc_1_Man_8-9_GlcNAc_2_ glycan (Fig. 1 *C*–*E*). Recruitment of MHC I with a mannose-only Man_9-7_GlcNAc_2_ glycan to the tapasin complex was not observed (Fig. 1 *C* and *D*). Notably, presence of the HLA-A*68:02-restricted, high-affinity peptide ETVSKQSNV (predicted affinity *K*_d_ ≈ 16 nM (72)) in the reaction mixture prevented handover of MHC I to the tapasin-ERp57 chaperone complex (*SI Appendix*, Fig. S2), reflecting selection and licensing of an MHC I charged with an optimal (immunogenic) peptide for cell surface presentation during MHC I quality control. Taken together, these findings indicate that TAPBPR-assisted reglucosylation of MHC I by UGGT1 governs the client exchange process.

### MHC I chaperone exchange is dependent on calreticulin

To determine if Crt is required for shuttling TAPBPR-chaperoned MHC I^glc.^ to tapasin in a concentration-dependent manner, we assessed the formation of complexes between monoglucosylated MHC I and the tapasin-ERp57-PaSta1 heterotrimer, both in the absence of Crt or in the presence of increasing concentrations of Crt (Fig. 2). When the tapasin-ERp57-PaSta1 heterotrimer (3 µM) was mixed with equimolar amount of monoglucosylated MHC I-TAPBPR in the absence of Crt, the SEC profile revealed two peaks corresponding to the individual complexes (Fig. 2 *A* and *B*). When Crt was added in substoichiometric amounts (1 µM), it primarily co-migrated with the tapasin-ERp57-PaSta1 heterotrimer and MHC I during SEC (Fig. 2 *A* and *B*). Compared to the tapasin-ERp57-PaSta1 heterotrimer alone, the peak shifted toward higher apparent molecular weight, indicating the formation of a pentameric complex comprising disulfide-linked tapasin-ERp57 with bound PaSta1 Fab, Crt, and MHC I. Providing a ten-fold molar excess of calreticulin (30 µM) relative to tapasin-ERp57-PaSta1 and MHC I did not enhance the formation of the pentameric complex. However, at this higher Crt concentration formation of a tetrameric tapasin-ERp57-Crt-PaSta1 complex apart from the pentameric complex was apparent (Fig. 2 *B*). These data show that the recruitment of calreticulin and MHC I^glc.^ to tapasin-ERp57-PaSta1 is dependent on the availability of monoglucosylated MHC I. Furthermore, we infer that either the dynamic binding equilibria between the MHC I quality control constituents are shifted towards the pentameric complex when calreticulin is present or that calreticulin actively withdraws monoglucosylated, peptide-receptive MHC I from TAPBPR. A further increase in calreticulin concentration leads to a tetrameric calreticulin-tapasin-ERp57-PaSta1 complex that deploys low-affinity interactions between calreticulin and the tapasin-ERp57 heterodimer.

**Fig. 2.**
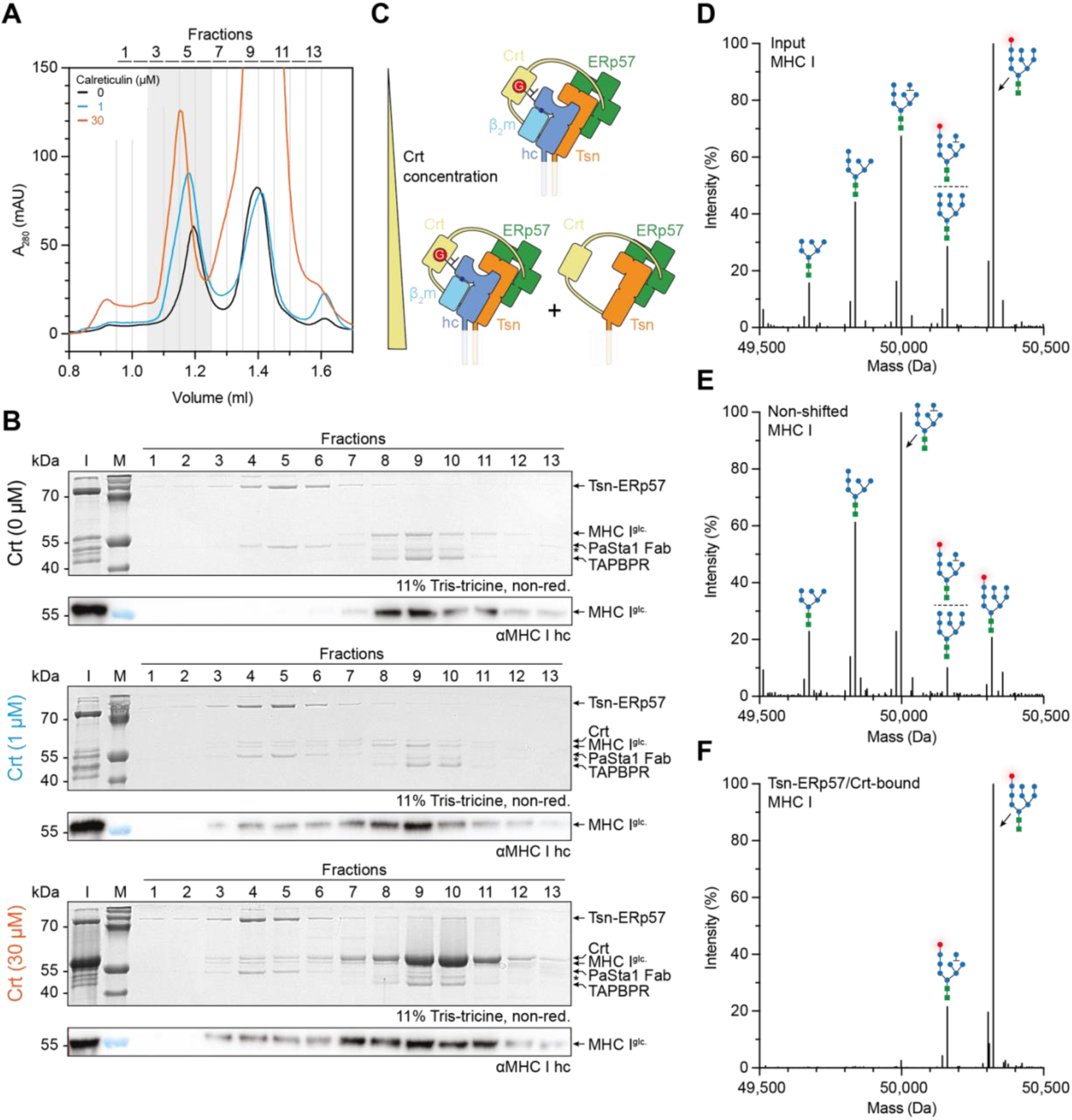
Calreticulin facilitates the transfer of monoglucosylated MHC I from TAPBPR to the tapasin-ERp57 quality control complex. (*A*) Size exclusion chromatography (SEC) profiles of monoglucosylated β_2_m-HLA-A*68:02/TAPBPR (3 µM) incubated with tapasin (Tsn)-ERp57-PaSta1 (3 µM) at varying concentrations of calreticulin (Crt): absence of Crt (0 µM), substoichiometric Crt (1 µM), and 10-fold access Crt (30 µM). Grey region indicates migration range of the Crt-MHC I-tapasin-ERp57-PaSta1 complex. (*B*) SDS-PAGE analyses of SEC fractions from (*A*) with immunoblot analysis probing for the MHC I heavy chain (αMHC I hc, HC10 antibody). (*C*) Schematic depiction of calreticulin concentration-dependent recruitment of monoglucosylated MHC I to the tapasin-ERp57 complex by Crt. (*D***–***F*) Deconvoluted mass spectra of single-chain β_2_m-HLA-A*68:02 before SEC (*D*, input MHC I), of the TAPBPR-associated β_2_m-HLA-A*68:02 (*E*, non-shifted MHC I). Peak labels show the structure of the N-glycan attached to single-chain β_2_m-HLA-A*68:02 (green dot – GlcNAc, blue dot – Man, red dot – Glc). Data presented in (*E*) relate to fraction 8 (1 μM Crt) in panel *B*, whereas data presented in (*F*) relate to fraction 4 (1 μM Crt) in panel *B*. M – marker, I – input, kDa – kilodalton, A_280_ – absorption at 280 nm, αMHC I – anti-MHC I heavy chain, * – impurity of β_2_m-HLA-A*68:02/TAPBPR complexes. Data in panels *A* and *B* are representative of three independent experiments (n = 3).

### Calreticulin facilitates the transfer of monoglucosylated MHC I to tapasin

The globular lectin domain of Crt specifically recognizes monoglucosylated glycans on its clients through a concave β-sheet. This β-sheet forms a long channel that establishes an extensive network of hydrogen bonds and hydrophobic interactions with the terminal four saccharides (Glc_1_Man_3_) of the glycan’s A-branch (73). The tetrasaccharide appears to be the minimal binding unit, exhibiting a low micromolar affinity (*K*_d_ of 0.7 µM) for Crt, which specifically recognizes the presence of the glucose moiety (73, 74). Using LC-MS, we assessed the molecular identity of the N-glycan of the shifted MHC I and compared it to the N-glycan of MHC I that was not transferred to the tapasin-ERp57-PaSta1 complex by Crt (Fig. 2 *D*–*F* and *SI Appendix*, Table S1). We observed that only monoglucosylated MHC I was recruited to the tapasin-ERp57-PaSta1 complex via Crt (Fig. 2 *F* and *SI Appendix*, Table S1), while MHC I with predominantly mannose-only glycans remained associated with TAPBPR during SEC (Fig. 2 *E* and *SI Appendix*, Table S1). These findings emphasize that the identity of the MHC I glycan is crucial for facilitating chaperone transfer corroborating results found in a cellular contexts (22).

### Calreticulin is recruited to TAPBPR-chaperoned, monoglucosylated MHC I

Peptide-receptive MHC I molecules are recruited to the PLC in association with Crt, where they receive and edit their peptide cargo (19, 20, 22). Empty, partially misfolded, or suboptimally loaded MHC I molecules that escaped initial ER quality control by the PLC are thought to be captured by TAPBPR and reglucosylated by UGGT1. These modified MHC I molecules were suggested to be re-engaged by calreticulin and directed back to the PLC for another round of peptide loading and editing (22–24). Thus, we hypothesized that MHC I-Crt intermediates must be detectable in our in-vitro handover experiment. To test if Crt can in a glycan-dependent manner extract MHC I from MHC I-TAPBPR complexes that are in dynamic equilibrium, we first mixed Crt and monoglucosylated, TAPBPR-chaperoned MHC I in stoichiometric ratios (3 µM each) and assayed changes in complex composition using a SEC-based assay (Fig. 3 *A*). When we compared the migration behavior of Crt in the Crt/monoglucosylated MHC I/TAPBPR mixture to Crt alone by SEC, we observed Crt and monoglucosylated MHC I co-migrating with larger hydrodynamic volume when compared to Crt alone (Fig. 3 *B* and *C*). Based on the glycan LC-MS analysis (Fig. 1 *B*, Fig. 2 *D*–*F*, and *SI Appendix*, Table S1), we assumed that monoglucosylated MHC I limited the amount of formed Crt-MHC I complex. To enhance complex formation, we increased the concentration of monoglucosylated, TAPBPR-chaperoned MHC I to 10 µM, which resulted in a higher abundance of the monoglucosylated MHC I-Crt complex, as detected by SDS-PAGE and immunoblot analysis (Fig. 3 *B* and *C*). In contrast, non-monoglucosylated MHC I did not induce a shift of Crt toward higher apparent molecular weight (Fig. 3 *C*). Notably, the Crt-monoglucosylated MHC I complex formed regardless of the presence or absence of high-affinity peptide (*SI Appendix*, Fig. S2 *B*, S3, and S4), indicating that our system can also capture Crt/MHC I complexes that have acquired optimal peptide during editing at the PLC. As expected, MHC I/TAPBPR complexes were dissociated in the presence of high-affinity peptides (*SI Appendix*, Fig. S3 *A* and *B*). Formation of Crt/monoglucosylated MHC I intermediates, as well as complexes with TAPBPR, was further validated by laser-induced liquid bead ion desorption (LILBID) mass spectrometry (MS) (Fig. 3 *D*). In summary, these data show that a monoglucosylated N-linked glycan of MHC I is required for Crt engagement and for withdrawal of MHC I from TAPBPR. Based on these findings, we propose that Crt actively extracts peptide-receptive MHC I from the TAPBPR-chaperoned state.

**Fig. 3.**
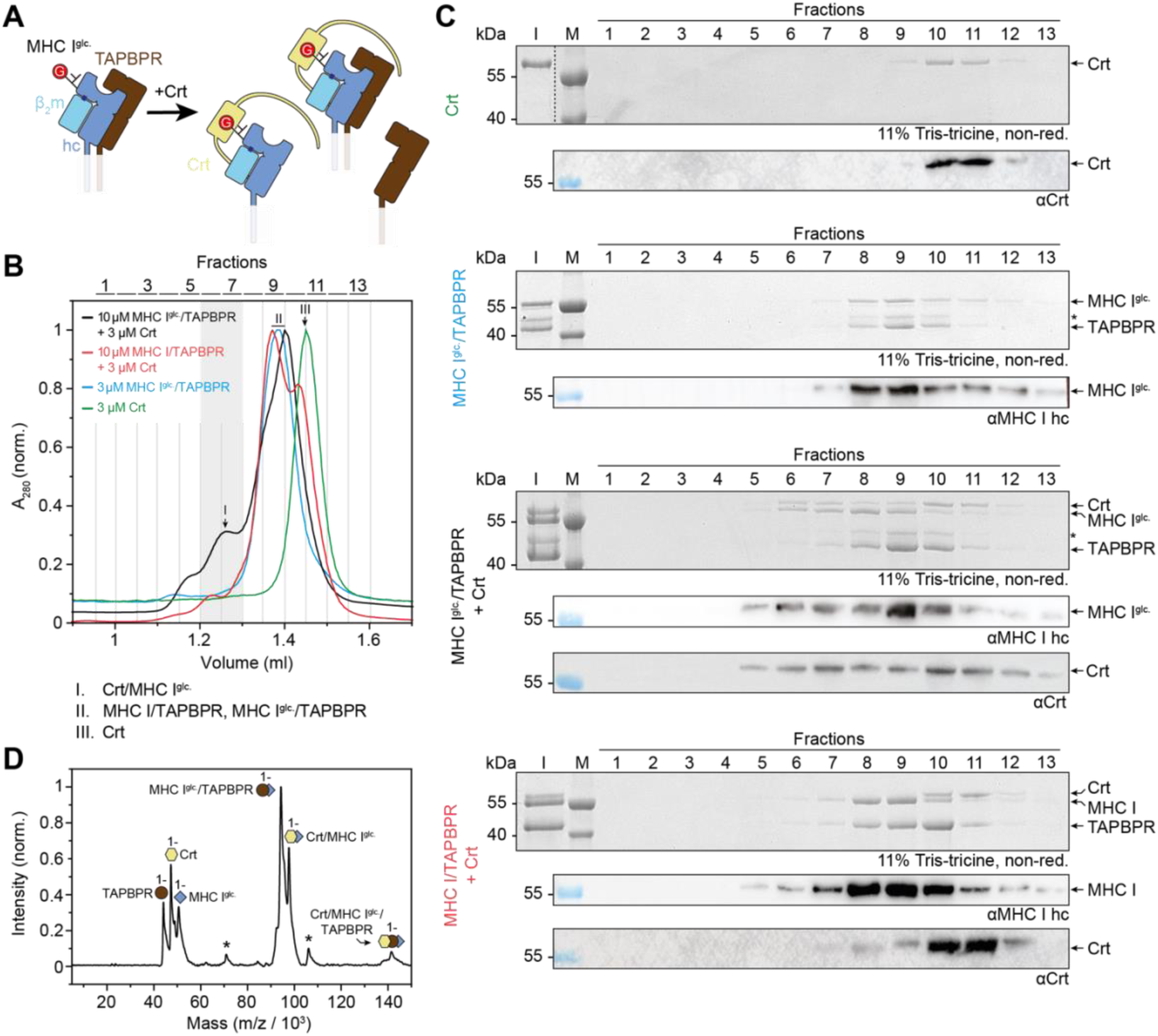
Monoglucosylation of MHC I attracts calreticulin to the TAPBR-chaperoned complex for chaperone exchange. (*A*) Schematic depiction of the formation of an MHC I intermediates upon calreticulin (Crt)-dependent engagement from the TAPBPR-chaperoned state. hc – MHC I heavy chain. (*B*) Size exclusion chromatography (SEC) profiles of Crt alone (3 µM, green line), monoglucosylated β_2_m-HLA-A*68:02/TAPBPR (3 µM, blue line), 3-fold molar excess of monoglucosylated β_2_m-HLA-A*68:02/TAPBPR (10 µM) over Crt (3 µM) (black line), and 3-fold molar excess of untreated β_2_m-HLA-A*68:02/TAPBPR (10 µM) over Crt (3 µM, red line). Grey region indicates migration range of the Crt/MHC I intermediate. Roman numbers above each peak relate to proteins or protein complexes defined below. (*C*) Non-reducing SDS-PAGE and immunoblot analysis of SEC fractions from (*B*). The HC10 antibody (αMHC I hc) was used to reveal the HLA-A*68:02 heavy chain. The FMC75 antibody (αCrt) was used to reveal Crt. (*D*) LILBID-MS analysis of formed complexes upon mixing Crt (3 µM) with the MHC I/TAPBPR complex (3 µM). Color coding is as in panel (*A*). Charge states of the individual proteins or complexes are indicated. Asterisks indicate unidentified complexes. M – marker, I – input, kDa – kilodalton, A_280_ – absorption at 280 nm, * – impurity of β_2_m-HLA-A*68:02/TAPBPR. Data in panels *B* and *C* are representative of three independent experiments (n = 3); data in panel *D* are representative of two independent experiments (n = 2).

### The acidic helix of calreticulin is dispensable for engaging monoglucosylated MHC I from TAPBPR

In the PLC, Crt interacts with the membrane-proximal immunoglobulin C1 (IgC1) domain of tapasin through an extended acidic helix (34, 60) that precedes the C-terminal KDEL ER-retention sequence. In a subset of myeloproliferative neoplasms, frameshift mutants of Crt have been identified, which are characterized by a change in the amino acid composition of their C-terminal domain and the absence of an ER-retention signal (75, 76), rendering the overall charge of the C-terminal helix positive. Frameshift mutants of Crt did not incorporate into the PLC and were predominantly secreted, likely due to the charge reversion and the absence of the ER retention signal (55). To explore the role of the C-terminal acidic helix in MHC I-chaperone exchange, we recombinantly produced a Crt variant, Crt^Δacidic^, lacking part of the acidic helix (Fig. 4 *A*). First, we investigated whether Crt^Δacidic^ could engage monoglucosylated MHC I in the MHC I/TAPBPR complex by comparing its effect on the monoglucosylated MHC I/TAPBPR complex to the mannose-only MHC I/TAPBPR complex (Fig. 4 *B*). Similar to wild-type Crt, a pronounced shift to higher apparent molecular weight appeared during SEC (Fig. 3 *B* and *C*, fractions 6 and 7), indicating co-migration of Crt^Δacidic^ and monoglucosylated MHC I. Co-migration of Crt^Δacidic^ with monoglucosylated MHC I was confirmed by SDS-PAGE and immunoblotting (Fig. 4 *C*). When we mixed MHC I that was not reglucosylated at its N-linked glycan with Crt or Crt^Δacidic^, we did not observe co-migration of MHC I with either Crt or Crt^Δacidic^ (Fig. 4 *B* and *C*). Thus, the C-terminal acidic helix of calreticulin is dispensable for engaging monoglucosylated MHC I in the TAPBPR-chaperoned state, while monoglucosylation of the MHC I glycan is essential for this process.

**Fig. 4.**
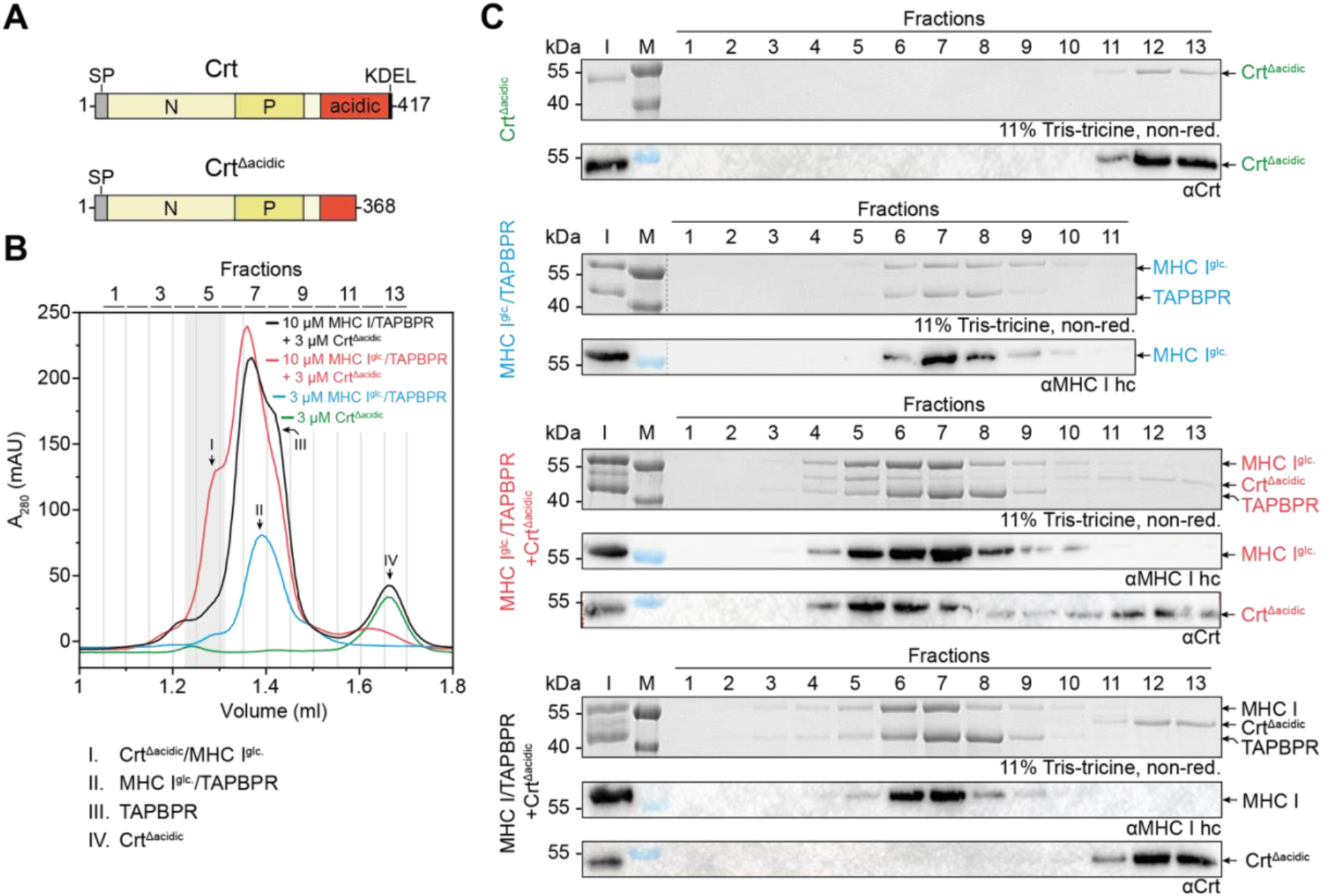
Calreticulin lacking its acidic helix binds monoglucosylated MHC I derived from a TAPBPR-chaperoned state. (*A*) Bar diagrams of wildtype calreticulin (Crt) and Crt with a truncated acidic helix (Crt^Δacidic^). (*B*) Size exclusion chromatography (SEC) profiles of Crt^Δacidic^ alone (3 µM, green line), monoglucosylated β_2_m-HLA-A*68:02/TAPBPR complex (3 µM, blue line), 3-fold molar excess of monoglucosylated β_2_m-HLA-A*68:02/TAPBPR (10 µM) over Crt^Δacidic^ (3 µM, red line), and 3-fold molar excess of untreated β_2_m-HLA-A*68:02/TAPBPR (10 µM) over Crt^Δacidic^ (3 µM, black line). Grey region indicates migration range of the Crt/MHC I intermediate. Roman numbers above each peak relate to proteins or protein complexes defined below. (*C*) Non-reducing SDS-PAGE and immunoblot analyses of MHC I heavy chain (αMHC I hc, HC10 antibody) or Crt (αCrt, FMC75 antibody) of SEC fractions shown in (*C*). KDEL – ER-retention signal lysine-aspartate-glutamate-leucine, SP – signal peptide N – N-domain; P – P-domain; A_280_ – absorption at 280 nm; kDa – kilodalton; I – input sample; M – protein size marker. Data in panels *B* and *C* are representative of three independent experiments (n = 3).

### The acidic helix of calreticulin is essential for chaperone exchange

Next, we examined whether the Crt^Δacidic^ variant facilitated transfer of MHC I from the TAPBPR-to the tapasin-chaperoned state. Crt engages in several low-affinity interactions with the editing module of the PLC (53). Aside from the lectin-Glc_1_Man_3_ interaction, calreticulin binds ERp57 via the tip of its P domain and cradles tapasin with its C-terminal acidic helix. These interactions collectively enable Crt to act as a “safety belt” for peptide-receptive MHC I at the PLC, ensuring tight structural association with tapasin (34, 60). Thus, we hypothesized that calreticulin mutants lacking one or more of these interactions are incapable of transferring monoglucosylated MHC I from TAPBPR to tapasin. After mixing the monoglucosylated MHC I-TAPBPR complex and tapasin-ERp57-PaSta1 with either wild-type Crt or its Crt^Δacidic^ variant, we observed distinct differences in the SEC profiles and SDS-PAGE analyses (Fig. 5). The presence of Crt, in combination with the monoglucosylated MHC I-TAPBPR complex and tapasin-ERp57-PaSta1, resulted in a shift of monoglucosylated MHC I to the tapasin-ERp57-PaSta1 complex migrating at higher apparent molecular weight (Fig. 5 *A*). On the other hand, the Crt variant lacking part of the C-terminal acidic helix was unable to transfer MHC I to tapasin-ERp57-PaSta1 as revealed by SEC and SDS-PAGE analysis (Fig. 5 *A* and *C*). Thus, the acidic helix of Crt is essential for docking MHC I to the tapasin-ERp57 complex. Notably, our in-vitro data are in line with results from cellular experiments showing that Crt frameshift mutants were unable to interact with the PLC and could not rescue peptide-receptive MHC I molecules (55).

**Fig. 5.**
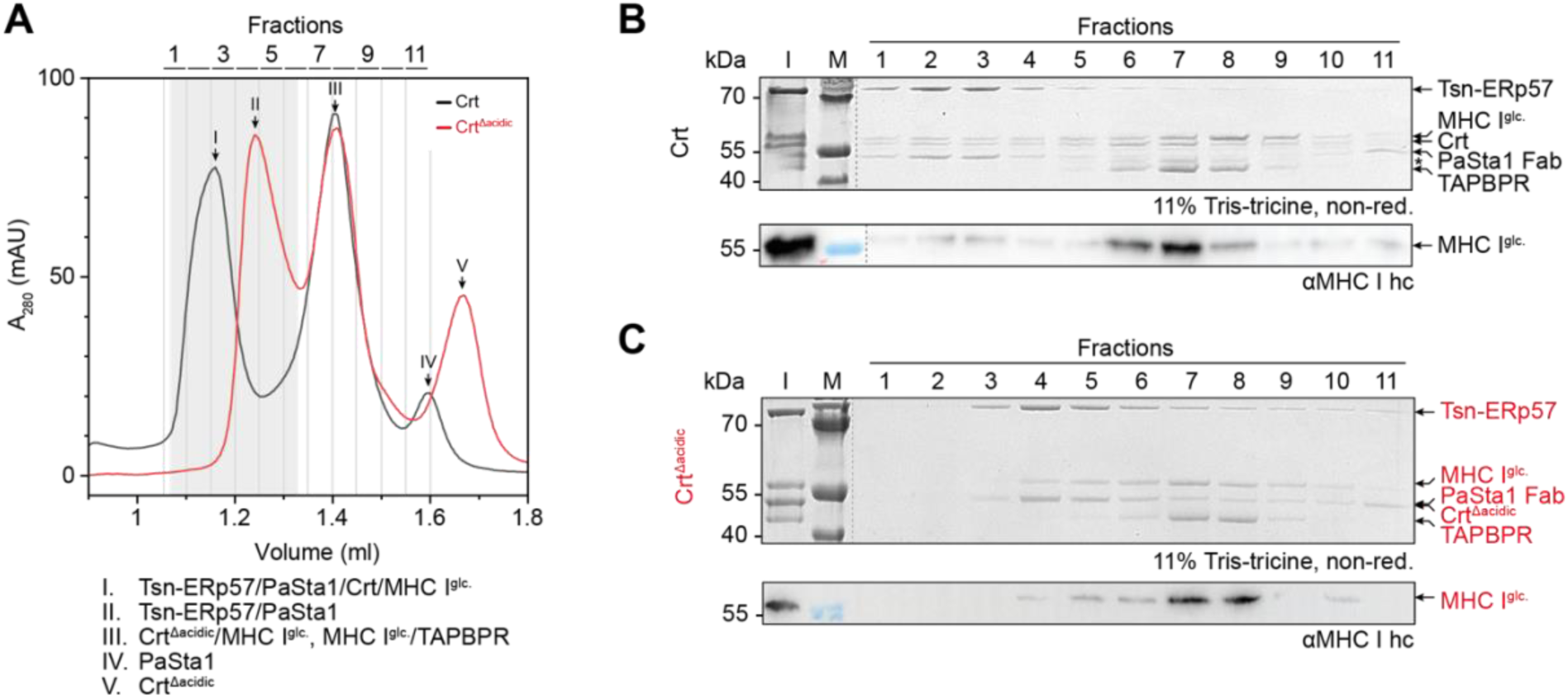
C-terminal acidic helix of calreticulin is essential for the exchange of monoglucosylated MHC I from TAPBPR to tapasin. (*A*) Size exclusion chromatography (SEC) profiles of calreticulin (Crt, black line) or Crt lacking the acidic helix (Crt^Δacidic^, red line) (3 µM each) incubated with the tapasin-ERp57-PaSta1 (3 µM) and monoglucosylated, peptide-receptive HLA-A*68:02/TAPBPR (3 µM). Grey region indicates migration range of the Crt-MHC I-tapasin-ERp57-PaSta1 and the Crt^Δacidic^-MHC I-tapasin-ERp57-PaSta1 complex. Roman numbers above each peak relate to proteins or protein complexes defined below. (*B*, *C*) Non-reducing SDS-PAGE and immunoblot analysis (αMHC I hc, HC10 antibody) of SEC fractions shown in (*A*), analyzing the ability of Crt (*B*) and Crt^Δacidic^ (*C*) to shift MHC I to tapasin-ERp57. A_280_ – absorption at 280 nm; kDa – kilodalton; I – input sample; M – protein size marker. Data in panels *A*, *B*, and *C* are representative of three independent experiments (n = 3).

## Discussion

In this study, we developed an in-vitro system with purified components to establish the chaperone exchange of MHC I from the secondary, downstream quality control machinery, composed of TAPBPR and UGGT1, to the primary quality control machinery, the PLC, in the ER. Central to this process is the reglucosylation of MHC I by UGGT1, which attaches a single glucose to the A branch of the N-core glycan, thus enabling engagement with the calnexin/calreticulin cycle and prolonging chaperone surveillance until proper folding is achieved. TAPBPR facilitates this process by bridging UGGT1 and immature MHC I, thus retaining it within the ER glycoprotein quality control cycle.

Our findings support a mechanistic model in which antigen presentation is controlled by a series of sequential quality control checkpoints that inspect MHC I complexes on their way to the cell surface. Consistent with previous reports that PLC-associated MHC I molecules carry exclusively monoglucosylated N-glycans (60), our results show that the concerted actions of UGGT1, TAPBPR, and calreticulin drive the retrograde trafficking of empty or misloaded MHC I molecules back to the PLC for additional rounds of peptide editing (Fig. 6). Although tapasin and TAPBPR exhibit preferences for different MHC I allomorphs (63, 77, 78), our results demonstrate that these systems can functionally cooperate to ensure high-fidelity antigen presentation.

**Fig. 6.**
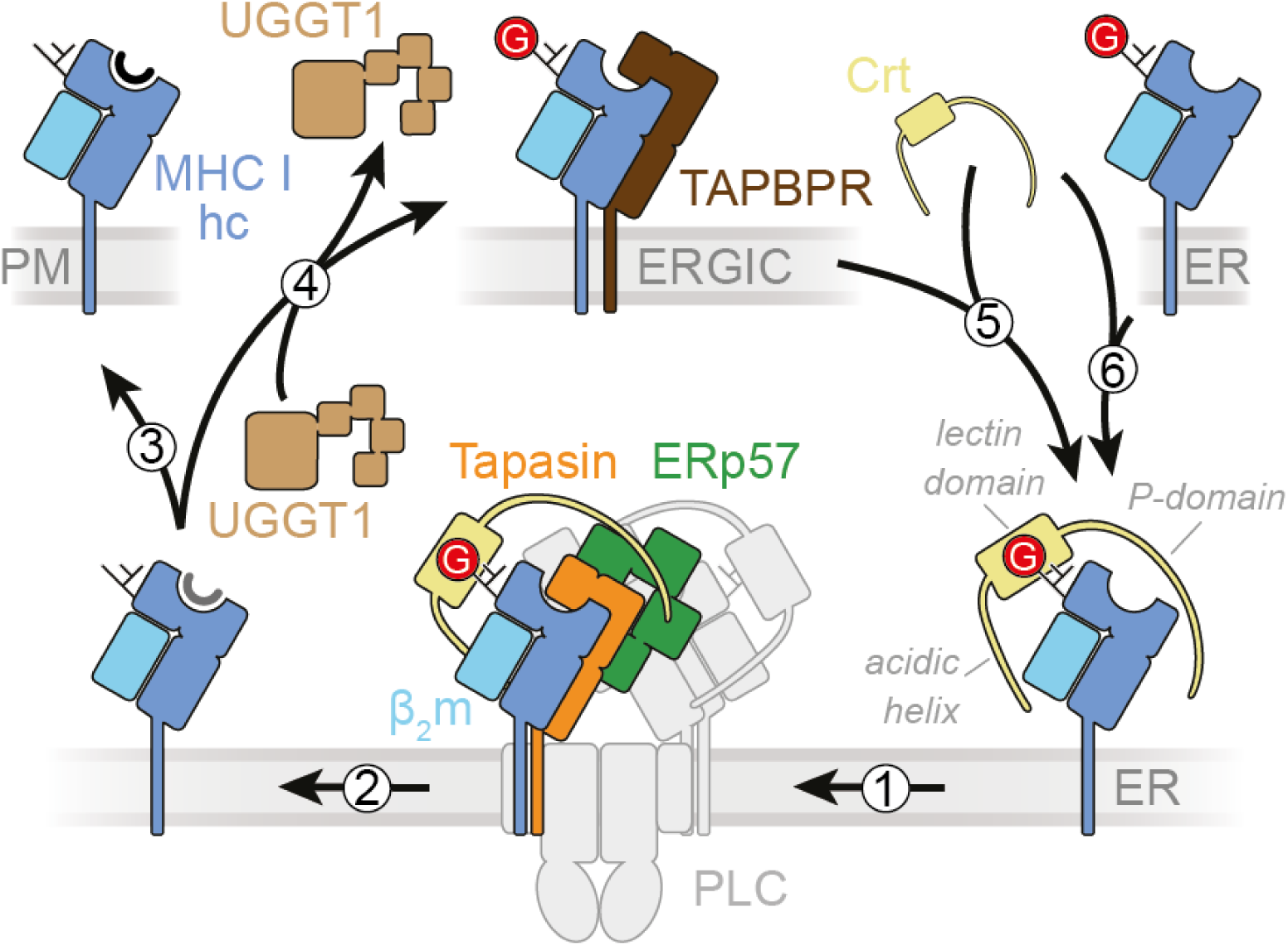
Model of MHC I recycling by a dedicated chaperone network. Monoglucosylated, peptide-receptive MHC I (MHC I hc, blue; β_2_m, light blue) is recognized and escorted by the lectin-like ER chaperone calreticulin (yellow) to the peptide-loading complex (PLC) for peptide loading and editing (1). MHC I molecules with optimal (high-affinity) peptide cargo are released from the PLC and fully deglucosylated by the action of α-glucosidase II that cleaves off the terminal glucose (2). MHC I with high-affinity peptides and a Man_9_GlcNAc_2_ glycan are licensed to traffic through the secretory pathway to present their antigenic cargo at the plasma membrane (PM) (3). Peptide-MHC I complexes that escaped the PLC with suboptimal (low-affinity) peptides are recognized in a secondary quality control step by TAPBPR and are reglucosylated by the glycoprotein folding sensor UDP-glucose:glycoprotein glucosyltransferase (UGGT1) (4). Reglucosylated peptide-receptive MHC I is engaged by calreticulin and trafficked back to the ER (5) where they can rejoin the PLC for another round of peptide loading and editing. The N-glycan of newly synthesized and translocated MHC I is trimmed by the action of glucosidases I and II to a monoglucosylated glycan which is recognized by calreticulin for docking to the PLC (6). Note that the extent to which different MHC I allomorphs depend either on tapasin or TAPBPR for editing of their peptide repertoire is also dictated by their intrinsic binding behaviors to the MHC I-specific chaperones.

At the PLC, calreticulin confines MHC I clients through three interaction sites involving its lectin domain, P domain, and C-terminal acidic helix that engage the MHC I glycan, ERp57, and tapasin, respectively (34, 60). Our experiments show that reglucosylation alone is sufficient for calreticulin to extract MHC I from TAPBPR. However, successful transfer to the tapasin-ERp57 editing module requires all three calreticulin interaction surfaces, indicating that fine-tuned dynamic equilibria between these specific, but low-affinity interactions are essential for effective chaperone exchange. Whereas calreticulin knockout cells still retain some MHC I-PLC association, these residual interactions may result from intrinsic MHC I-tapasin affinity or compensatory roles played by calnexin (61). However, the extent to which calnexin contributes to accompany newly synthesized MHC I to the PLC or retrieving escaped MHC I from post-ER compartments back to the PLC remains to be elucidated. Concomitant release of Crt and MHC I from the PLC following peptide loading, however, agrees with the observation that Crt does not associate with the PLC in the absence of MHC I (22, 79).

Additionally, our study sheds light on one aspect of functional consequences of calreticulin frameshift mutants associated with myeloproliferative neoplasms (55, 76). These mutations, characterized by loss of the ER-retention signal (KDEL) and charge reversal at the C-terminal helix (55, 76), exhibit impaired association with the PLC and diminished MHC I surface expression. Although our data do not explain the transformative potential of the Crt frame-shift mutant (80), our in-vitro system mimics the PLC-centric phenotype by failing to promote MHC I transfer to the tapasin-ERp57 complex (55).

Importantly, we identified a stable intermediate complex of calreticulin and peptide-receptive MHC I during the handover process. The existence of this intermediate demonstrates that calreticulin alone can act as a specific chaperone for MHC I, driven by the concerted action of its lectin and P domain. These biochemically tractable intermediates help stabilize inherently unstable, peptide-free MHC I molecules, not only before their engagement by tapasin-ERp57, but also during their retrograde trafficking from TAPBPR back to the PLC.

Given that MHC I allomorphs can be classified into tapasin-dependent and –independent ones based on their surface expression in tapasin-deficient cells (63, 78), we focused our study on the well-characterized, TAPBPR-dependent HLA-A*68:02 allomorph (64). Interestingly, in our assay, HLA-A*68:02 transferred efficiently from TAPBPR to a tapasin-chaperoned state, suggesting that both tapasin-dependent and tapasin-independent MHC I allomorphs can pass through the PLC to acquire high-affinity antigenic peptides.

We hypothesize that TAPBPR-mediated MHC I quality control and subsequent retrograde trafficking of MHC I represent a prominent mechanism of MHC I quality control past the PLC. Alternative pathways for retrograde trafficking, such as passive diffusion or calnexin-assisted transport, lack the capacity to discriminate between MHC I loaded with high-affinity peptide and those that are peptide-receptive or misloaded with low-affinity peptides. Observations of MHC I interacting with UGGT1 in absence of TAPBPR (24) likely reflect recognition of partially unfolded MHC I molecules rather than a mechanism for fine-tuning the peptide repertoire. Consistent with this notion, loss of the association between TAPBPR and UGGT1 reduced the quality control exerted on the MHC I allomorph HLA-A*68:02 (24). As recognition of MHC I molecules by UGGT1 can occur in a TAPBPR-independent or in a TAPBPR-dependent manner (22), the extent to which each of the two pathways are preferred is dictated by the molecular identity and structural states of a given MHC I allomorph. Thus, monoglucosylation by UGGT1 and subsequent binding of Crt to unfavorable MHC I molecules, including semi-stable, peptide-receptive MHC I, MHC I bound to non-optimal peptides, or MHC I heavy chains that lack β_2_m, seems to be the overarching principle for retrograde trafficking of MHC I to the PLC (24).

Two categories of MHC I molecules have been introduced to describe TAPBPR-dependencies of MHC I allomorphs (81): (i) generalists, which bind a broad and diverse peptide repertoire with lower specificity, and (ii) specialists, which present a narrow, high-specificity peptide repertoire. Generalist MHC I allomorphs are intrinsically less stable due to the structural flexibility required to accommodate a broad peptide repertoire, with peptide binding serving as a key stabilizing factor. Specialists typically achieve immediate stabilization upon binding one of their few high-affinity peptides. Thus, generalists more often acquire moderate-or low-affinity peptides, rendering them more reliant on TAPBPR-mediated chaperoning and redirection to the PLC for optimal peptide editing. The limited peptide exchange efficiency of TAPBPR in post-ER compartments across different MHC I allomorphs (82), combined with alterations in the peptide repertoire observed in the presence of the C94A TAPBPR mutant, which disrupts UGGT1-mediated reglucosylation (24), further supports a primary role for TAPBPR in peptide–MHC I surveillance and recycling.

### Limitations of the Work

While our in-vitro system offers precise control and high mechanistic resolution for dissecting MHC I recycling, it inevitably simplifies the complex and dynamic environment of the ER. Our focus on a single TAPBPR-dependent MHC I allomorph (HLA-A*68:02) – chosen for its strong TAPBPR affinity and biochemical tractability – restricts the generalizability of our findings. MHC I allomorphs with weaker TAPBPR affinity or different peptide-binding affinities may rely differently on reglucosylation and calreticulin-mediated retrieval.

Moreover, our system excludes potential modulatory contributions from other ER-resident factors, post-ER quality control pathways, and membrane-associated processes that influence antigen presentation in vivo. Consequently, while our system enables detailed dissection of core chaperone interactions and handover steps, the absence of membrane context, cofactors, and spatial compartmentalization may alter the kinetic equilibria and affinity profiles of MHC I-chaperone interactions in an in-vivo context.

## Materials and Methods

### Plasmid constructs

Single-chain β_2_m-HLA-A*68:02 (HLA-A68), human TAPBPR, human tapasin^ΔTMD^-ERp57^C36A^, and *Ct*UGGT were previously described (17, 48, 65). PaSta1 Fab constructs (heavy and light chain) were kindly provided by David H. Margulies (49). Human, full-length calreticulin (UniProtKB P27797) with a N-terminal, TEV-cleavable His_6_-tag lacking the signal peptide (amino acids 18-417) and the calreticulin variant lacking part of the C-terminal acidic helix (amino acids 18-369) were cloned into the pETM-11 vector, respectively.

### Cell culture and transfection of HEK293-F/Expi293 cells

HEK293-F or Expi293 cells were grown in FreeStyle 293 Expression Medium (Gibco) at 37 °C, 5% CO_2_, and shaken at 120 rpm. For coexpression of the MHC I^Fos^ and TAPBPR^Jun^, 5 ml expression medium was supplemented with 75 μg of β_2_m-HLA-A68^Fos^ and 75 μg of TAPBPR^Jun^ DNA (equimolar ratio) and mixed with a four-fold molar excess of linear PEI (Polysciences) diluted in 5 ml expression medium. Similarly, coexpression of PaSta1 Fab was performed with a 1:2 molar ratio of PaSta1 Fab DNA of heavy chain (50 µg) and light chain (100 µg). In accordance, expression of *Chaetomium thermophilum* (*Ct*) UGGT was performed with 150 µg of *Ct*UGGT plasmid DNA. All plasmid DNA used for transfection was purified using the PureYield Plasmid Midiprep System (Promega). After incubation for 30 min, the transfection mixture was added to 50 ml of HEK293-F cells at a density of 16×10^6^ cells/ml. Three hours post transfection, cells were diluted to 3×10^6^ cells/ml and 3.5 mM valproic acid (Merck Millipore), and 10 μM kifunensine (Biomol) were added. The culture supernatant containing the secreted proteins was harvested five days after transfection.

### Purification of *Ct*UGGT

To purify secreted *Ct*UGGT from transfected HEK293 cells, the culture supernatant was cleared from cells by centrifugation at 2,000×g for 10 min and dialyzed twice against 1xPBS (1.5 mM KH_2_PO_4_, 8 mM Na_2_HPO_4_, 137 mM NaCl, 2.7 mM KCl, pH 7.6) for 3 h and overnight, respectively. Ni^2+^-NTA beads (Thermo Fisher Scientific) were added to the supernatant and stirred over night at 4 °C. Metal affinity chromatography was performed in PBS, supplemented with 20 mM imidazole and 5% (v/v) glycerol as wash buffer. *Ct*UGGT was eluted with 1xPBS supplemented with 200 mM imidazole and 5% (v/v) glycerol.

Monodisperse UGGT was isolated by size-exclusion chromatography (SEC, Superdex 200 Increase 10/300, Cytiva) in 1xHBS buffer (50 mM HEPES-NaOH pH 7.4, 150 mM NaCl). Protein fractions were concentrated by ultrafiltration (Amicon Ultra, 30-50 kDa MWCO, Merck) and stored at –80 °C.

### Purification of MHC I^Fos^/TAPBPR^Jun^ complexes

HEK293 culture was cleared from cells by centrifugation at 2,000×g for 10 min. Avidin (IBA Lifesciences, 7.5 mg/l) and Tris-HCl pH 8.0 (25 mM final) was added to the cleared supernatant. Affinity chromatography was performed with Strep-Tactin Sepharose (1 ml/l) (IBA Lifesciences) using 1xTBS buffer (25 mM Tris-HCl pH 8.0, 100 mM NaCl). The tethered complex was eluted in 1xTBS supplemented with 5 mM desthiobiotin. Leucine zippers were removed by overnight digestion at 4 °C with Tobacco Etch Virus (TEV) protease. MHC I/TAPBR complexes were isolated either by size exclusion chromatography (SEC) on a Superdex 200 Increase 10/300 column (Cytiva) in 1xHBS buffer and concentrated by ultrafiltration (Amicon Ultra 50 kDa MWCO, Merck) or subjected to UGGT-mediated reglucosylation before SEC purification.

### UGGT-mediated reglucosylation

MHC I-TAPBPR complexes (3 μM) were incubated with *Ct*UGGT (1 μM) in 1×HBS buffer supplemented with 200 μM UDP-glucose (UDP-Glc) and 5 mM CaCl_2_ for 10 min at RT. The reaction was stopped by adding 25 mM of EGTA and MHC I-TAPBPR complexes were purified via SEC (Superdex 200 10/300 Increase, Cytiva) equilibrated in 1xHBS buffer. MHC I-TAPBPR peak fractions were pooled, flash-frozen in liquid nitrogen, and stored at – 80 °C. Samples of the purified MHC I-TAPBPR complex were analyzed by LC-MS.

### Protein production of calreticulin variants

Crt and Crt^Δacidic^ were produced in *Escherichia coli* Rosetta(DE3)pLysS cells. In brief, cells were transformed with plasmids encoding Crt and Crt^Δacidic^. Overnight cultures of 10–20 ml LB media (10 g tryptone, 5 g yeast extract, 10 g NaCl per 1 l) supplemented with kanamycin were used to inoculate 2 L of LB main culture. The main culture was incubated at 37 °C and 180 rpm until an optical density at 600 nm (OD_600_) of 0.4–0.6 was reached. Temperature was reduced to 20 °C and protein production was induced by addition of 1 mM isopropyl-β-D-thiogalactopyranoside (IPTG). After 20 h, cells were harvested by centrifugation at 4 °C and 4,500×g for 15 min. Cells were flash-frozen in liquid nitrogen and stored at –80 °C.

### Protein production in insect cells

The disulfide-linked tapasin-ERp57 heterodimer was produced in *Spodoptera frugiperda* (*Sf*21) cells as described (48). Cells were grown in *Sf*900 II SFM medium (Thermo Fisher Scientific) at 28 °C and transfected with modified EMBacY bacterial artificial chromosomes (BACs) using X-tremeGENE DNA transfection reagent (Roche). After incubation for 72 h at 28 °C, recombinant baculovirus V0 was harvested and utilized for production of amplified baculovirus V1. The tapasin-ERp57 heterodimer was expressed in 800 ml suspension culture at a cell density of 10^6^ cells/ml by infection with 0.5–2.0% (v/v) baculovirus V1. Cells were harvested 72 h post cell proliferation arrest by centrifugation at 4 °C and 1,300×g for 10 min. Cell pellets were flash-frozen in liquid nitrogen and stored at –80 °C.

### Purification of tapasin-ERp57 heterodimers

Infected *Sf*21 cell pellets were resuspended in 100 mL lysis buffer (50 mM HEPES-NaOH pH 7.4, 150 mM NaCl, 25 mM imidazole, 1 mM phenylmethylsulfonyl fluoride (PMSF), 1 mM benzamidine) per 1 l expression culture and lysed by sonication. After centrifugation at 20,000×g, 4 °C for 30 min, the supernatant was incubated with pre-equilibrated Ni^2+^-*N*-nitrilotriacetic acid (NTA) agarose resin (Thermo Fisher Scientific) for 1 h at 4 °C. The resin was washed with lysis buffer without protease inhibitors and bound proteins were eluted with elution buffer (1xHBS supplemented with 300 mM of imidazole). Protein fractions were pooled, the His_6_-tag was cleaved, and the buffer was exchanged to 1xHBS buffer by dialysis over night at 4 °C. Tapasin-ERp57 heterodimers were concentrated by ultrafiltration (Amicon Ultra 50 kDa, MWCO, Merck) and isolated by SEC (Superdex 200 10/300 Increase, Cytiva) equilibrated in 1xHBS buffer. Peak fractions were pooled, flash-frozen in liquid nitrogen, and stored at –80 °C.

### Purification of calreticulin variants

Cell pellets of *E. coli* Rosetta(DE3)pLysS containing expressed Crt^WT^ and Crt^Δacidic^, respectively, were resuspended in 100 ml lysis buffer (50 mM HEPES-NaOH pH 7.4, 150 mM NaCl, 25 mM imidazole, 5 mM CaCl_2_, 1 mM phenylmethylsulfonyl fluoride (PMSF), 1 mM benzamidine, Benzonase 1:10,000 (v:v) from Merck Millipore) per 1 l expression culture and lysed by sonication. The lysate was centrifuged for 30 min at 20,000×g, 4 °C and the supernatant incubated with pre-equilibrated Ni^2+^-NTA agarose resin (Thermo Scientific) for 1 h at 4 °C. The Ni^2+^-NTA agarose resin was washed with lysis buffer without protease inhibitors and Benzonase. The immobilized protein was eluted with elution buffer (1xHBS supplemented with 300 mM of imidazole and 1 mM CaCl^2^). Protein fractions were pooled, the His_6_-tag was cleaved, and the buffer was exchanged to 1xHBS buffer supplemented with 1 mM CaCl_2_ by dialysis over night at 4 °C. Crt^WT^ or Crt^Δacidic^, respectively, were concentrated by ultrafiltration (Amicon Ultra 10 kDa, MWCO, Merck) and isolated by SEC (Superdex 200 10/300 Increase, Cytiva) equilibrated in 1xHBS buffer supplemented with 1 mM CaCl_2_. Peak fractions were pooled, flash-frozen in liquid nitrogen, and stored at –80 °C.

### Purification of PaSta1 antibody fragment

Secreted PaSta1 Fab was purified from cleared HEK293 culture supernatant after centrifugation at 2,000×g, 4 °C for 10 min. The cleared supernatant of HEK293 F was supplemented with Tris-HCl pH 8.0 (25 mM final) and affinity chromatography was performed with Ni^2+^-NTA agarose resin (Thermo Scientific). The resin was washed with 1xHBS buffer supplemented with 25 mM imidazole and bound protein was eluted with 1xHBS supplemented with 300 mM imidazole. Protein fractions were pooled and the buffer was exchanged over night by dialysis against 1xHBS buffer at 4 °C. PaSta1 Fab was isolated by size-exclusion chromatography (SEC, Superdex 200 Increase 10/300, Cytiva) in 1xHBS buffer. Protein fractions were concentrated by ultrafiltration (Amicon Ultra, 10 kDa MWCO, Merck) and stored at –80 °C.

### Complex formation of various MHC I-chaperone complexes

SEC analyses of each component of the chaperone exchange reaction were performed with 3 µM of each protein used in the experiment, unless stated otherwise. Tapasin-ERp57 was pre-incubated with PaSta1 Fab in equimolar ratio for at least 15 min 4 °C prior to experiments. Prior to exchange experiments involving peptide-loaded HLA-A*68:02, the MHC I complex was incubated over night at 4 °C with 250x molar excess of the high-affinity peptide ETVSKQSNV (predicted affinity of 16 nM (83)). Exchange experiments were performed by incubation of several exchange components for 30 min at 4 °C. SEC analyses were performed with a Superdex 200 3.2/300 Increase column (Cytiva) equilibrated with 1xHBS at a flow rate of 0.05 ml/min. At least 20 µl samples of each SEC fraction were gathered and analyzed by SDS-PAGE and immunoblotting.

### Liquid chromatography (LC)-MS analysis

LC-MS analyses were conducted on a BioAccord System (Waters). Top-down LC-MS data for glycan analysis was acquired using a cone voltage of 45 V, 1.5 kV capillary voltage, and a desolvation temperature of 500 °C on an ACQUITY UPLC Protein BEH C4 Column, 300 Å,

1.7 μm, 2.1×50 mm or 150 mm (Waters) at 80 °C applying a linear water/acetonitrile gradient (5–45% (v/v) acetonitrile in 8.0 min or 5–95% (v/v) acetonitrile in 66.5 min) supplemented with 0.1% (v/v) formic acid. Mass spectra were recorded in positive polarity at 2 Hz in full scan mode at 400–7000 m/z. Reglucosylation mixtures were spun down at 16,000×g for 10 min and directly analyzed by LC-MS. Glycoprotein masses were calculated and confirmed using the software platform UNIFY 3.1.0 (Waters). Combined intact mass spectra of glycoproteins were deconvoluted in UNIFY 3.1.0, utilizing the quantitative MaxEnt1 algorithm iterating to convergence with 1.0 Da model width. Deconvoluted spectra were centroidized based on peak height and plotted in OriginPro 2022b (Origin Lab Corporation, USA).

### Laser induced liquid bead ion desorption (LILBID)-MS analysis

Samples were prepared for native LILBID-MS analysis by performing a buffer exchange into 200 mM ammonium acetate, using Zeba Spin desalting columns (Thermo Fisher Scientific, USA). The samples, at concentrations of 3 to 10 µM, were loaded into the piezo-driven droplet generator (MD-K-130, Microdrop Technologies GmbH, Germany) operating at 10 Hz and generating 50 µm diameter droplets. Sample droplets were irradiated by an IR laser (a Nd:YAG pumped optical parametric oscillator (OPO)) with a wavelength of 2.8 µm in a high-vacuum irradiation chamber. A laser pulse of 6 ns pulse length was used to cause explosive expansion of the sample droplets leading to the release of ionized molecules into a homebuilt time-of-flight analyzer by applying an acceleration voltage (2.6 kV). The procedure was performed with Wiley McLaren type ion optics, by setting both first and second lens to –4 kV and then ramping the first lens 5 to 25 µs after the explosion to –6.6 kV for 370 µs. A reflectron, working at –7.2 kV, guides the ions into the Daly type detector, optimized for high m/z. Data processing was performed by using OriginPro 2022b (Origin Lab Corporation, USA).

### SDS-PAGE and immunoblotting

6× sodium dodecyl sulfate (SDS)-loading buffer (150 mM Tris-HCl pH 6.8, 30% (v/v) glycerol, 1.2% (w/v) SDS, 0.02% (w/v) Bromophenol Blue) was added to SDS-PAGE samples and incubated at 95 °C for 10 min. 10 μl of samples were loaded on 11% Tris-tricine gel and gel electrophoresis was performed at 180 V for 45 min. The gels were either stained with InstantBlue Coomassie Protein Stain (abcam) or blotted onto a PVDF membrane (0.45 µm) by semi-dry blotting for 30 min at 25 V. The blotted membranes were blocked in 5% (w/v) milk powder in TBS-T for 1 h and incubated with an appropriate primary antibody at 4 °C overnight. MHC I hc was analyzed with the anti-HLA-A,B,C primary antibody HC10 (in-house production, hybridoma, mouse, 1:20 dilution) and Crt^WT^/Crt^Δacidic^ with the anti-Crt primary antibody FMC75 (abcam, mouse, 1:2,000 dilution). Blots were washed three times in TBS-T at RT for 10 min each and then incubated with a corresponding second antibody-HRP conjugate (anti-mouse, A-2554 from Sigma-Aldrich, goat, 1:20,000) for 1 h at RT. After three times wash with TBS-T for 10 min each at RT, membranes were incubated with Clarity Western ECL reagent (BioRad).

## Data availability

Source data are provided at https://doi.org/10.25716/gude.18f9-c6gc.

## Acknowledgments and funding sources

We acknowledge the support by the European Research Council (ERC Advanced Grant 789121 to R.T. and ERC Advanced Grant 101141396 to R.T.), the German Research Foundation (DFG Grant TA157/12-1 to R.T. and DFG Grant 557111829 to N.M.) and the Collaborative Research Center grant (CRC1507/P18 to R.T. and P13 to N.M.). We thank Dr. Lina Sagert and Dr. Ines K. Müller for support and helpful advice. We are grateful to all members of the institute for discussion and Inga Nold and Andrea Pott for editing of the manuscript. We are grateful to Dr. David H. Margulies for providing plasmids encoding PaSta1 Fab light chain and heavy chain.

